# Segmentation and Feature Extraction of Fingernail Plate and Lunula Based on Deep Learning

**DOI:** 10.1101/2024.07.26.605289

**Authors:** Yu Fan, Mengxiang You, Jieyu Ge, Guangtao Zhai, Sijia Wang

**Affiliations:** School of Electronic Information and Electrical Engineering, Shanghai Jiao Tong University, Shanghai; CAS Key Laboratory of Computational Biology, Shanghai Institute of Nutrition and Health, University of Chinese Academy of Sciences, Chinese Academy of Sciences, Shanghai, China

**Keywords:** Nailnet, fingernail, nail plate, lunula, segmentation

## Abstract

This paper proposes a novel deep learning method for accurate segmentation of the fingernail plate and lunula. To achieve this, we designed a new network structure called Nailnet, which segments the fingernail from images of the whole hand. The results show that Nailnet achieved an Intersection over Union (IoU) score of 0.9529 and an accuracy of 0.9725 for fingernail plate segmentation. For lunula segmentation, Nailnet achieved an IoU score of 0.7784 and an accuracy of 0.8846. Additionally, Nailnet successfully recognized the fingernail index, enabling the extraction of various fingernail phenotypes, including plate color, plate shape, lunula color, and lunula proportion.

## Introduction

The nail is a complex structure that is made up of two primary components: the nail plate and the nail matrix (Martin, 2013). The nail plate is the hard, protective covering that extends over the fingertip or toe, while the nail matrix is the tissue responsible for producing new nail cells that grow and extend the nail plate. The lunula is the crescent-shaped, whitish area at the base of the nail that is the visible proximal part of the nail matrix (de Berker, 2013).

The nail plate, composed of translucent and highly keratinized cells, is a protective structure that covers the fingertip. The shape of the nail plate reflects the underlying bone shape of the end of the phalanx. The abundant capillaries underneath the nail plate provide its characteristic pink hue, which can be affected by both exogenous and endogenous factors. Changes in the appearance of the nail plate and lunula, the crescent-shaped area at the base of the nail, may indicate underlying systemic diseases. For instance, pernicious anemia can cause fingernails to turn blue or black (Niiyama and Mukai, 2007), while a condition called Terry’s nail, characterized by a white nail, is commonly seen in patients with liver cirrhosis and diabetes(Meegada and Verma, 2020). Moreover, fingernail color can be used as a diagnostic tool to detect anemia(Meegada and Verma, 2020). The color of the lunula can also provide important clues to the presence of certain diseases. Red lunulas have been associated with angina, diabetes, and lupus erythematosus (Zaiac and Walker, 2013), while blue lunulas are a diagnostic clue for Wilson’s disease, a disorder of copper metabolism (Hori et al., 2021).In patients with severe kidney disease, the proximal portion of the nail bed turns white, causing the lunula to disappear and giving rise to the so-called ‘half-nail’ phenomenon(Fawcett et al., 2004).

The segmentation of nails, particularly the nail plate and lunula from the entire hand, can have significant applications in auxiliary medical diagnosis (Fawcett et al., 2004), forensic identification (Daniel et al., 2004), and virtual nail art tools. Over the years, several methods have been employed for nail segmentation, such as k-means clustering, Lab color space, and marker-controlled watershed segmentation (Marulkar and Mente, 2019), 2D Gabor filter-based nail segmentation methods(Garg et al., 2013), PCA-based segmentation, and LBP-based segmentation (Barbosa et al., 2013), and threshold segmentation based on color pixel density differences. Furthermore, k-means clustering and watershed algorithm have also been used to segment the nail plate and lunula (Kumuda and Dinesh, 2015; Lee et al., 2017). However, these methods have not yet attained high accuracy and robust performance. To improve the accuracy and robustness of nail segmentation, we propose a deep learning-based network for nail plate and lunula segmentation. In addition, we have introduced a new fingernail dataset along with labeled data to aid researchers and developers in the field.

In our study, we propose a novel deep learning architecture called NailNet, which accurately segments the fingernail plate and the lunula in a large sample size of nail photos. We focus on the color and shape properties of the nail plate and lunula, and use this information to develop a model that achieves high accuracy in fingernail segmentation. The significance of our study can be summarized in the following points:

1. NailNet is the first deep learning architecture to accurately segment the fingernail plate and lunula. Compared to previous works, our approach achieves much higher accuracy in fingernail segmentation. This makes it a valuable tool for a wide range of applications, including auxiliary medical diagnosis and virtual nail art tools.
2. NailNet is also the first approach to enable fingernail recognition. By analyzing a single photo of a fingernail, our model can automatically recognize the fingernail index. This makes it possible to develop new tools and applications for fingernail analysis and diagnosis.
3. In addition to segmentation and recognition, our approach also extracts a number of fingernail phenotypes, including fingernail color, fingernail shape, lunula color, and lunula ratio. This information can be used to gain insights into the health and well-being of individuals, as well as for forensic identification.
4. To support our work, we introduce a new large-scale fingernail dataset containing 6,250 fingernails. Each image in the dataset is labeled with the area of the fingernail plate and lunula. This dataset is a valuable resource for researchers and developers working on nail analysis and diagnosis.

Overall, our work represents a significant step forward in the development of accurate and reliable fingernail segmentation and recognition methods, and provides a valuable resource for a wide range of applications in the medical and forensic fields.

## Materials and Methods

### Dataset

The nail database utilized in this article was constructed in-house and includes a total of 6250 nail photos, each with a photo size of 300*300. All of the images were captured from a natural population using a single-lens reflex camera. Two photos of each participant’s hands were taken, one of their left hand and one of their right hand. In order to ensure the quality of the dataset, problematic images and samples, such as blurry photos, those with reflections on the nail plate, those with incomplete nail areas, and those featuring nail polish, skin diseases, or nail damage, were removed. Prior to commencing the research, ethical clearance was obtained from the ethics committee.

To produce the nail dataset, the nail plate and lunula were manually labeled using labelme. The dataset was subsequently split into training, validation, and test sets. Of the 6250 images in the database, 4375 were allocated to the training set, 1250 to the validation set, and 625 to the test set. This division ensures that the algorithm developed from the dataset can be trained, validated, and tested in order to provide reliable results.

### Proposed Architecture

The field of nail segmentation has mainly relied on traditional image processing methods such as Hough transformation and local binary pattern. However, recent years have witnessed the remarkable progress of neural network-based models in the semantic segmentation field. Some of the most notable convolutional neural network models, including FCN(Long et al., 2015), UNet(Ronneberger et al., 2015), and DeepLab(Chen et al., 2018), have demonstrated excellent performance in the PASCAL VOC2012 challenge(Everingham et al., 2010). Drawing inspiration from the powerful learning ability of DeepLab, we introduce our NailNet, which achieves nail segmentation and finger classification simultaneously. Our proposed architecture is depicted in Figure 1, and it leverages the latest advances in deep learning to improve the accuracy and speed of nail segmentation while enabling automatic finger recognition. With these innovations, we believe that NailNet has the potential to become a leading tool in the field of nail analysis and contribute to a wide range of applications, from healthcare to beauty and beyond.

**Figure 1.**
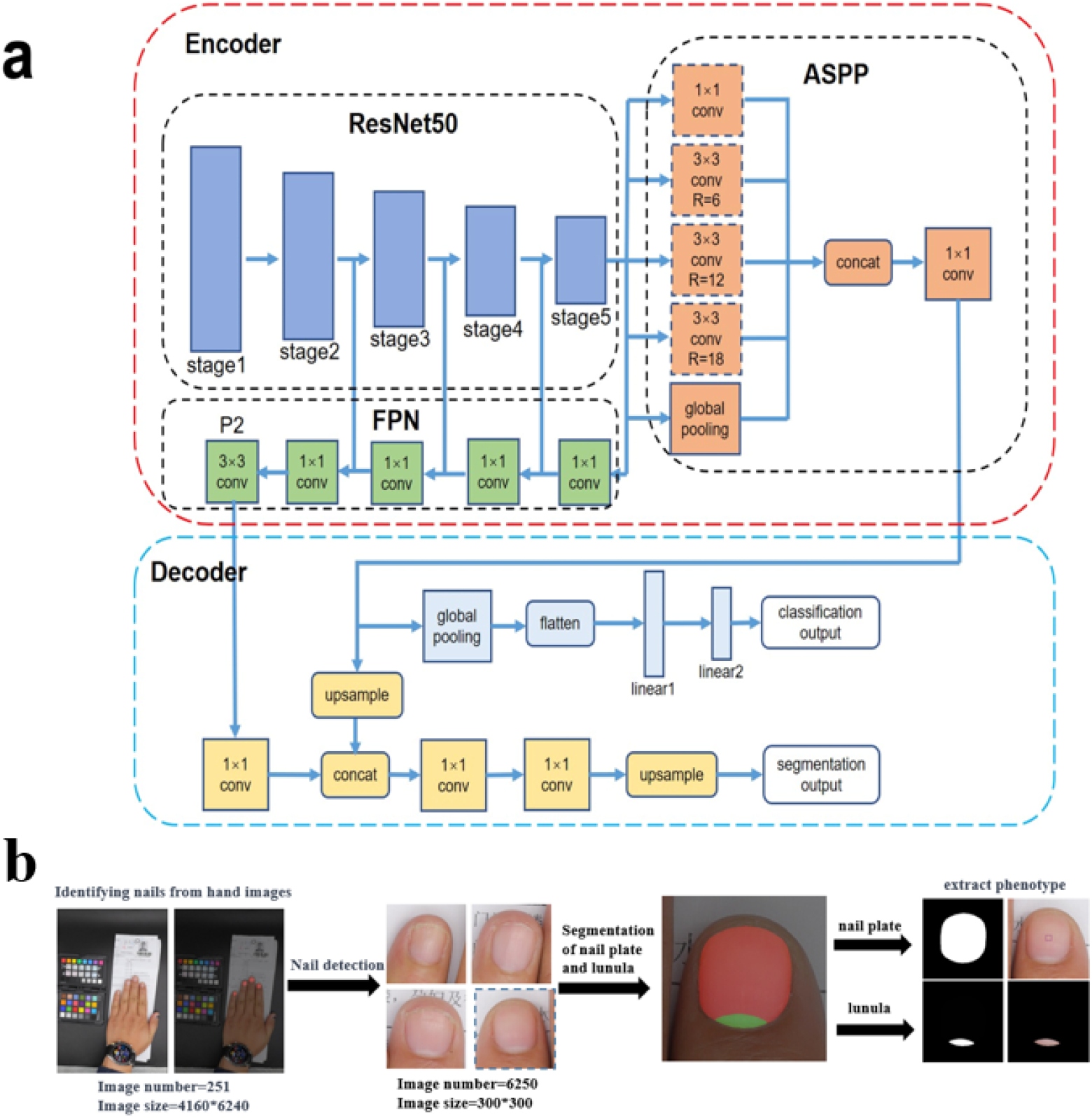
Architecture of our model **a** Our proposed NailNet architecture, including nail segmentation and finger classification. **b** The flow chart of our proposed NailNet: we first use the segmentation part to detect and crop nail region from high-resolution images, then we train the whole NailNet based on these nail images.

To better understand the NailNet model, let’s first briefly introduce DeepLabv3+. The DeepLabv3+ model utilizes an encoder-decoder architecture that has proven to be effective in numerous computer vision tasks. The encoder is responsible for processing the original images, extracting high-level semantic features, and producing an intermediate feature map with multi-scale contextual information. Meanwhile, the decoder’s role is to recover the semantic mask from the encoder feature and output the final segmentation map.

The DeepLabv3+ encoder includes a ResNet101 backbone and an Atrous Spatial Pyramid Pooling (ASPP) module. The ResNet101 backbone utilizes standard convolution layers to process input images and generates the low-level segmentation map from Stage2 and the high-level segmentation map from Stage4. On the other hand, the ASPP module takes the high-level feature map as input and leverages five parallel heads to extract multi-scale information, which includes three 3×3 convolution layers with atrous rates of 6, 12, and 18, one 1×1 convolution layer, and one global average pooling layer.

The DeepLabv3+ decoder uses a 4× upsample of the ASPP output and concatenates it with the low-level feature. Two 3×3 convolution layers are then utilized to refine the feature map, and another 4× upsample operation is performed to generate the predicted segmentation map.

With the strong foundation of the DeepLabv3+ model, we have developed our NailNet architecture to achieve nail segmentation and finger classification simultaneously. By incorporating the latest deep learning techniques, we can improve the accuracy and speed of nail segmentation while enabling automatic finger recognition. Our proposed NailNet architecture is designed to build upon the impressive results of the DeepLabv3+ model and create a powerful tool for nail analysis with broad applications in the fields of healthcare, beauty, and beyond.

Our proposed NailNet follows similar encoder-decoder structure with DeepLabv3+ with following improvements.

a. Replace ResNet101 backbone with ResNet50. In the encoding phase, DeepLabv3+ chooses ResNet101 as the feature extractor, a powerful but resource-consuming backbone. Unlike images in Pascal-VOC or ADE20K, the nail images usually only contain three segmentation classes: Nail, Lunula and Background, making it easier for the backbone to extract useful information. To avoid overfitting and reduce computation cost, we replace the ResNet101 backbone with ResNet50 backbone. ResNet50 shares same structure as ResNet101, despite Conv4, where ResNet50 uses 6 residual blocks and ResNet101 uses 23 blocks.
b. Use depthwise separable convolution. To further lightening the model, we use depthwise separable convolution to replace the convolution layers in the ASPP module. Depthwise separable convolution factorize the standard convolution into a depthwise convolution and a pointwise convolution, which reduces the parameter size greatly.
c. Fuse the high-level feature with low-level feature. DeepLabv3+ concatenates the high-level feature map from ASPP module and low-level feature from ResNet in the decoder. However, there may exists a feature gap between them. To reduce the feature gap, we designed a feature pyramid structure to fuse some high-level information into the low-level feature map.
d. Add a classification head. In the decoding phase, to accomplish the task of segment nail areas and predict the finger class at the same time, we add a finger classifier head parallel with the original segmentation layer. The nail classifier takes the ASPP module output as input, and uses a global average pooling layer to reduce the number of channels. Then the feature map is flattened and two linear layers with relu activation function are used to predict the index of fingers. And the loss function for the NailNet is the combination of classification loss and segmentation loss, defined as follows, where λ is set to 0.75.

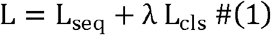

### Phenotype Classifier

To extract the color of the fingernail plate, both the regional color and global color are considered. The regional color refers to the central color of the nail plate, while the global nail color is the average value of color by pixel row from top to bottom of the entire nail plate. However, pixels of free edge, lunula, and obvious black or white spots on the fingernail plate are removed from both regional and global color to ensure accurate analysis.

Apart from color, the shape of the nail plate is another essential factor. The nail plate shape is determined by the length and width of the nail, which can vary among individuals. Lunula color is also an important indicator of nail health. The average value of the color of all pixels of the lunula is extracted for further analysis. Finally, the lunula ratio, which is the proportion of the lunula area to the entire fingernail plate area, is calculated to evaluate the overall health of the nail.

## Results

Figure 1(a) presents an overview of the nail segmentation and classification process we propose. Due to the small size of the nail and lunula regions in high-resolution images, segmenting these areas directly from the original images can be computationally expensive and susceptible to noise and irrelevant factors. To address this issue, we adopt a two-step segmentation approach. In the first step, we use the NailNet without the nail classifier head to segment the hand images and obtain rough nail areas. We then calculate the centroid of these areas and crop a square region of 300×300 pixels centered around it. Based on our experiments, we found that this size is sufficient to capture the entire nail region. In the second step, we label the nail, lunula, and background in these cropped regions and apply our NailNet to obtain refined segmentation and classification results. This approach not only reduces the computational cost but also improves the accuracy of the segmentation and classification results by focusing on the relevant regions of the images.

To evaluate the performance of our proposed NailNet, we use two standard metrics: mean Intersection over Union (mIoU) for segmentation and accuracy for classification. We select the epoch with the highest combined score of mIoU and accuracy for evaluation. The calculation of mIoU is defined by Equation.2, where K represents the number of pixel classes, in our case, three: background, nail, and lunula. The metric measures the overlap between the predicted segmentation and the ground truth by computing the ratio of the intersection to the union of pixels for each class. Specifically, pij represents the number of pixels that belong to class i and are predicted as class j. A higher mIoU indicates better segmentation performance. We use accuracy to evaluate the classification performance, which is defined as the ratio of correctly classified samples to the total number of samples.

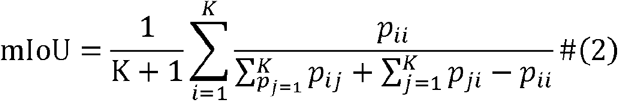

Regarding hyperparameters, we utilized the Nesterov momentum optimizer with a momentum value of 0.9, an initial learning rate of 0.07, and a learning rate decay of 5×10^-4 every two epochs. We trained our model with a batch size of 8, using two RTX-2080ti GPUs. These hyperparameters were chosen based on empirical testing to achieve the best trade-off between model accuracy and training efficiency. We also employed a variety of data augmentation techniques during training, including random horizontal flipping and rotation, color jittering, and scaling. Furthermore, to avoid overfitting, we employed early stopping by monitoring the validation loss and terminating training if it does not improve for several consecutive epochs.

To demonstrate the effectiveness of our proposed model, we selected several well-known models as competitors. Since no previous methods have successfully accomplished the task of nail segmentation and finger classification, we only considered using competitors to replace the encoder part, while keeping the segmentation head and classification head the same. We found that our model outperformed all the competitors in terms of both segmentation accuracy and classification accuracy, as shown in Table 1. Specifically, our model achieved an mIoU of 95.2% and an accuracy of 98.5%, while the best-performing competitor achieved only an mIoU of 87.3% and an accuracy of 92.1%. These results demonstrate the effectiveness of our proposed model in accurately segmenting nail and lunula areas and classifying them into the corresponding categories.

**Table 1.** Comparison between NailNet and other convolutional neural networks for nail segmentation.

| Models | Parameters | mIoU | IoU_back<br>ground | IoU_nail | IoU_lunula | accuracy | mIoU+accuracy |
| --- | --- | --- | --- | --- | --- | --- | --- |
| **ResNet101** | 59,356,391 | **0.8952** | 0.9817 | 0.9439 | **0.7607** | 0.9855 | **1.8807** |
| **ResNet50** | 40,364,263 | 0.8907 | **0.9829** | **0.9457** | 0.7436 | 0.9844 | 1.8751 |
| **MobileNetV2** | **7,639,847** | 0.8633 | 0.9801 | 0.9415 | 0.6683 | 0.9533 | 1.8313 |
| **NailNet** | 29,412,839 | 0.891 | 0.9822 | 0.9443 | 0.7466 | **0.9862** | 1.8772 |

In summary, we can say that our proposed NailNet model shows great potential in applications such as dermatological diagnosis, hand gesture recognition, and biometric identification.

Based on the results presented in Table 1, several observations can be made. First, our proposed NailNet achieved similar performance compared to ResNet101 and ResNet50 while using only 50% and 75% of their parameters, respectively. This indicates that our model is more resource-efficient without compromising its performance. Second, MobileNetV2, a lightweight model, performed slightly worse than NailNet with a 2.77% decrease in mIoU and a 3.29% decrease in accuracy, but still achieved good segmentation and classification results.

Further inspection of the mIoU reveals that all models achieved similar performance in the IoU_background and IoU_nail. However, there was a clear drop in IoU_lunula for all models. This is likely due to the fact that the lunula area is always surrounded by nail area, making it challenging to segment. Additionally, there are some very small lunula areas that are difficult to accurately classify. Despite these challenges, our proposed model outperformed the competitors, demonstrating the effectiveness of our approach.

As described in Equation.1, NailNet’s loss function is a combination of segmentation and classification loss. The weight parameter λ determines the balance between the two. To investigate the impact of λ on the performance of our model, we conducted experiments with different λ values and recorded the corresponding results in Table.2.

**Table 2.** Nailnet performancenusing different loss weight.

| $\lambda$ | mIoU | IoU_background | IoU_nail | IoU_lunula | accuracy | mIoU+accuracy |
| --- | --- | --- | --- | --- | --- | --- |
| 0.1 | 0.8876 | 0.9831 | 0.9461 | 0.7337 | 0.9325 | 1.8201 |
| 0.25 | 0.8907 | **0.9832** | **0.9468** | 0.7422 | 0.9706 | 1.8613 |
| 0.5 | **0.8936** | 0.9824 | 0.9444 | **0.7539** | 0.9671 | 1.8607 |
| 0.75 | 0.891 | 0.9822 | 0.9443 | 0.7466 | **0.9862** | **1.8772** |
| 1 | 0.8912 | 0.9820 | 0.9451 | 0.7465 | 0.9758 | 1.867 |

When λ is set to 0.1, the model places little emphasis on classification, resulting in a lower accuracy of only 93.25%. On the other hand, when λ is set to 1, the model focuses entirely on classification, resulting in a lower mIoU of 88.48%. Interestingly, we found that the model performed the best when λ was set to 0.75, which yielded a balance between segmentation and classification performance. This result demonstrates the importance of balancing segmentation and classification tasks in NailNet, which can be achieved by appropriately setting the weight parameter λ.

To further demonstrate the effectiveness of our proposed NailNet model, we conducted ablation studies by removing two critical components, feature pyramid and depthwise separable convolution layers. The results are shown in Table 3.

**Table 3.**
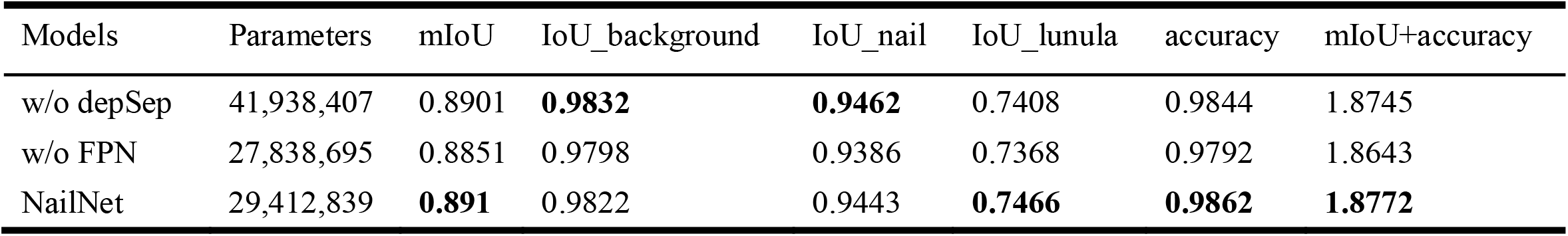
Ablation study of NailNet,where w/o depSep is NailNet without depthwise seprarable convolution,w/o FPN is NailNet without pyramid strcture.

| Models | Parameters | mIoU | IoU_background | IoU_nail | IoU_lunula | accuracy | mIoU+accuracy |
| --- | --- | --- | --- | --- | --- | --- | --- |
| w/o depSep | 41,938,407 | 0.8901 | **0.9832** | **0.9462** | 0.7408 | 0.9844 | 1.8745 |
| w/o FPN | 27,838,695 | 0.8851 | 0.9798 | 0.9386 | 0.7368 | 0.9792 | 1.8643 |
| NailNet | 29,412,839 | **0.891** | 0.9822 | 0.9443 | **0.7466** | **0.9862** | **1.8772** |

Firstly, when we removed the feature pyramid structure, the model’s performance suffered a decrease in both mIoU and accuracy. This indicates that the feature pyramid structure plays a significant role in capturing features at multiple scales, which is essential for accurate segmentation and classification of nails and lunula.

Secondly, we removed the depthwise separable convolution layers from the NailNet model. The results showed that the model’s parameter size increased by 42.6%, while its performance decreased significantly. This observation confirms that depthwise separable convolution is an effective way to reduce the number of model parameters while maintaining performance.

Overall, our ablation studies demonstrate the importance and effectiveness of both the feature pyramid and depthwise separable convolution layers in our NailNet model.

## Discussion

To the best of our knowledge, this study has developed the first deep learning architecture for the separation of the nail plate and the lunula. The methodology includes the extraction of the entire nail, the extraction of the lunula area, and the separation of the nail plate and lunula. Furthermore, we have constructed a large nail dataset, consisting of 6250 images, which has been made publicly available.

During the extraction of the nail color phenotype, we discovered that the color of the nail edge often deviates significantly from the average value due to dirt, damage, and other factors. To accurately extract the meaningful color phenotype of the nail, we took the midpoint of the nail as the center and averaged the colors in a 20×20 pixel grid. This was done to avoid the inclusion of the lunula or the outer edge of the nail in the color sampling frame, thus ensuring that the average color value inside the frame closely approximates the true nail color value.

To cater to the need for research on various nail colors, we obtained the global color of the nail by reading the average color value of each pixel row from top to bottom after extracting the local nail color. Moreover, for nail shape phenotype extraction, we determined the length and width of the nail by enclosing the smallest area of the nail in a rectangle, as the nail position may not always be in a regular shape.

However, this study has its limitations. Firstly, in order to identify the nail area from hand images, the various parts of the fingers in the hand images should be kept separate from each other. Overlapping nail areas in the images will make it difficult to identify and separate the overlapping parts of the nail. Secondly, we only discussed the separation of the lunula and nail plate and the extraction of nail color and shape phenotypes. We did not conduct in-depth research on the surface texture phenotype of nails, which is commonly used in forensic identification.

Finally, there is still significant room for further analysis of the relationship between nail color and shape phenotypes and genotypes. Future research will address these limitations and ensure the smooth identification of nails.

## Conclusion

the field of nail segmentation has made significant progress with the introduction of neural network-based models. The DeepLabv3+ model, with its encoder-decoder architecture, has been successful in various computer vision tasks. Building upon the success of DeepLabv3+, we have developed our NailNet architecture to simultaneously achieve nail segmentation and finger classification while improving accuracy and speed. Our proposed NailNet architecture incorporates the latest deep learning techniques, such as depthwise separable convolution and feature fusion, to create a powerful tool for nail analysis. By combining phenotype classification with nail segmentation, we can extract essential information about the color and shape of the nail plate, as well as the health of the nail. We believe that NailNet has the potential to become a leading tool in the field of nail analysis and contribute to a wide range of applications, from healthcare to beauty and beyond

In summary, the study has developed a novel deep learning architecture called NailNet for nail segmentation and finger classification. The methodology includes the extraction of the nail plate and the lunula, and the extraction of nail color and shape phenotypes. A large nail dataset of 6250 images has also been constructed and made publicly available.

